# Dental caries in human evolution: frequency of carious lesions in South African fossil hominins

**DOI:** 10.1101/597385

**Authors:** Ian Towle, Joel D. Irish, Isabelle De Groote, Christianne Fernée

**Author notes:** **Contact details:** Ian Towle, Room 352, James Parsons Building, Byrom Street, Liverpool, L3 3AF.

## Abstract

Caries frequencies in South African fossil hominins were observed and compared with other hominin samples. Species studied include *Paranthropus robustus*, *Homo naledi*, *Australopithecus africanus*, *early Homo* and *A. sediba*. Teeth were viewed macroscopically with Micro-CT scans used to confirm lesions. Position and severity of each lesion were also noted and described. For all South African fossil hominin specimens studied, 16 have carious lesions, six of which are described for the first time in this study. These are from a minimum of six individuals, and include four *P. robustus,* one *H. naledi,* and one *early Homo* individual. No carious lesions were found on deciduous teeth, or any teeth assigned to *A. africanus*. Most are located interproximal, and only posterior teeth are affected. Caries frequency typically ranges between 1-5% of teeth in non-agricultural human samples, and this pattern seemingly holds true for at least the past two million years in the hominin lineage. Non-agricultural populations significantly above or below this threshold generally have a specialized diet, supporting other dietary evidence that *A. africanus* likely consumed large amounts of tough, non-cariogenic vegetation. Given the common occurrence of caries in the other hominin species, cariogenic bacteria and foods were evidently common in their collective oral environment. Along with recent research highlighting additional examples of caries in *H. neanderthalensis*, early *Homo* and Pleistocene *H. sapiens*, caries is clearly an ancient disease that was much more common than once maintained throughout the course of human evolution.

## Introduction

Caries can form when specific intra-oral bacteria demineralize dental tissue through the release of acids as they metabolize sugars and starches (Larsen et al., 1991; Byun et al., 2004). Different bacteria may be involved, and the genome of one in particular, *Streptococcus mutans*, suggests it has evolved quickly in the last few thousand years, potentially in response to human population growth and agriculture (Nishikawara et al., 2007; Cornejo et al., 2013). For active lesions to form the oral pH must be lowered locally over extensive periods (Gussy et al., 2006). Therefore, frequency and location vary with diet and behavior. Some foods are especially cariogenic, with those containing high levels of refined carbohydrates and sugars particularly virulent (Clarkson et al., 1987; Prowse et al., 2008; Rohnbogner & Lewis, 2016). Many fruits, honey, and some nuts and seeds can also be cariogenic (Humphrey et al., 2014; Novak, 2015). Tough and fibrous foods are linked with low caries rates, since they tend to create a more alkaline oral environment from high levels of saliva production (Moynihan, 2000; Prowse et al., 2008; Rohnbogner & Lewis, 2016). Meat and certain plants have also been associated with low caries frequency (Moynihan, 2000; Novak, 2015).

Environmental and genetic influences are also important to consider when researching caries prevalence. The clearest example is that of moderate levels of fluoride in drinking water, which decrease the likelihood of carious lesions developing (Kotecha et al., 2012; Slade et al., 2013). It is not yet clear how genetic difference between populations influence caries rate (e.g., Haworth et al., 2018), although, differences in dental morphology between hominin groups will likely affect the position and frequency of caries, particularly when comparing distantly related groups. However, the teeth, dental tissues, and crown positions that are affected in a population can give further insight into the diet, the oral microbiome, and food processing behaviors (Kelley et al., 1991; Shen et al., 2004; Meinl et al., 2010; Bignozzi et al., 2014; Novak, 2015; Takahashi & Nyvad, 2016).

Caries is widely reported in archaeological samples, with rates varying substantially in agricultural groups (e.g., Lunt, 1974; Varrela, 1991; Slaus et al., 1997; Watt et al., 1997; Srejic, 2001; Vodanovic et al., 2005; Caglar et al., 2007; Esclassan et al., 2009; Slaus et al., 2011; Vodanovic et al., 2012; Novak, 2015). A wide variety of living and fossil species also show evidence of caries, including dinosaurs, fishes, bats, bears, and primates (Palamra et al., 1981; Lovell, 1990; Trinkaus et al., 2000; Kear, 2001; Kemp, 2003; Miles & Grigson, 2003; Sala et al., 2007; Lanfranco & Eggers, 2012; Lacy, 2014; Arnaud et al., 2016). In the hominin lineage it is often suggested that caries is a modern disease and is scarce or absent in past hominin groups, usually justified by inferring dietary or oral bacterial differences between present day and ancient hominin populations (Brothwell, 1963; Armelagos & Cohen, 1984; Hildebolt & Molnar, 1991; Tillier et al., 1995; Lanfranco & Eggers, 2012; Guatelli-Steinberg, 2016; Adler et al., 2017). However, evidence for the presence of carious lesions in a variety of non-agricultural hominin groups is growing (e.g., Grine et al., 1990; Trinkaus et al., 2000; Lanfranco & Eggers, 2012; Lacy et al., 2012; Humphrey et al., 2014; Lacy, 2014; Liu et al., 2015; Arnaud et al., 2016; Margvelashvili et al., 2016).

In light of this evidence, South African fossil hominin material was re-analyzed, and *H. naledi* recorded for the first time, for the presence of caries; comparisons were then made with other fossil hominins and recent human samples. Nine carious lesions have already been recorded in the South African hominin fossils, including two in a mandible belonging to an early *Homo* individual, SK 15 (Clement, 1956). The rest are attributed to *Paranthropus robustus*, with three on SKX 5023 (Grine et al., 1990), two lesions on SK 55 (Robinson, 1952), one on SK 13/14 (Robinson, 1952), and one on DNH 40 from Drimolen (Moggi-Cecchi et al., 2010; Towle et al., 2019). We hypothesize that caries is likely more common than currently thought, and utilizing micro tomography (micro-CT) scans may help in getting a better understanding of the frequency of lesions in different species. Given substantial dietary differences have been suggested between the species studied (e.g., Kupczik et al., 2018; Towle et al., 2017; Henry et al., 2012; Peterson et al., 2018), we expect there will be significant differences in caries rate. The results may therefore give further information into dietary changes in the hominin lineage.

## Materials and methods

The material studied includes specimens assigned to Early *Homo*, *Australopithecus sediba*, *Homo naledi*, *A. africanus* and *Paranthropus robustus*. A full list of the specimens are available in Towle (2017). All material studied is curated at the University of the Witwatersrand and The Ditsong National Museum of Natural History. Only complete teeth were included in analysis. Each tooth was examined macroscopically under good lighting with a 10x hand lens used for initial classification of lesions. The frequency of caries for each samples is calculated as follows:

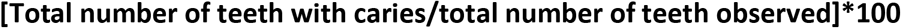

A carious lesion was recorded when there is a clear cavitation; color changes alone were not diagnostic. The severity and position of lesions on a tooth were also recorded. Caries severity was scored on a scale of 1 to 4 following Connell and Rauxloh (2003), with (1) enamel destruction only; (2) involvement of dentine but pulp chamber not exposed; (3) destruction of dentine with the pulp chamber exposed; (4) gross destruction with the crown mostly destroyed. Lesion location is also recorded as: distal, buccal, occlusal, lingual, root, mesial and gross, or a combination of these.

The interaction between caries and other dental pathologies is often complex. It has been suggested that because caries can lead to Antemortem Tooth Loss (ATML), correction methods need to be implemented (e.g., Kelley et al., 1991; Lukacs, 1995; Duyar & Erdal, 2003). However, caries correction methods are not appropriate to implement in this research. Although the majority of missing (i.e., extracted) teeth today are due to caries, in many past populations this would not have been the case. Severe attrition and fractures that exposed the pulp, and periodontal disease, are all other possible contributors. In particular, given the high rates of wear in fossil hominin specimens, it is likely that most cases of ATML resulted from attrition, not caries. Therefore, following Meinl et al. (2010) and Larsen et al. (1991) no corrective methods were implemented. By not including correction methods direct comparisons with other populations can be made. Additionally, maxilla and mandible fragments are so rare in the fossil record that to include ATML data would have little effect on overall frequencies.

Micro-CT scans of particular teeth are included in this study to help clarify if a cavity is carious, and to show the extent of lesions. Micro-CT scans can differentiate between normal enamel and dentine, and areas affected by caries, due to the lower density of areas affected with this pathology (Swain & Xue, 2009; Neves et al., 2010; Boca et al., 2017). Tertiary dentine formation can also be associated with carious lesions, and can be observed on Micro-CT scans (Towle, 2019; Towle et al., 2019). Using such techniques can clearly make visible the extent of a lesion, even if the cavity on the surface is ambiguous (Rossi et al., 2004; McErlain et al., 2004). Scans were completed by the Department of Human Evolution, Max Planck Institute for Evolutionary Anthropology with a BIR ACTIS 225/300 (kV, 100 mA, 0.25 brass filter) or a SkyScan 1172 (100 kV, 94 mA, 2.0 mm aluminium and copper filter) micro-CT scanner. The isometric voxel sizes resulting from these scans range between 15 and 50 micrometres (mm) (M. Skinner, personal communication, 2018).

## Results

In addition to the six carious teeth already described in the literature for South African fossil hominins, an additional six are added in this study (Table 1). From the six newly identified teeth, four are from *P. robustus* specimens and two from *H. naledi*. In total there is now a minimum of 16 carious lesions across the South African fossil hominin material (Table 2). The four new *P. robustus* teeth come from two individuals. Therefore, there are four *P. robustus* individuals, one *H. naledi* individual, and one *early Homo* individual that exhibit caries. No carious lesions were found in deciduous teeth, or any tooth belonging to *A. africanus* and *A. sediba*. In sum, there are 16 carious lesions on 12 teeth belonging to six individuals so far described in the South African fossil hominin material. Table 1 does not include the carious *P. robustus* tooth from *Drimolen* (DNH 40), as this sample was not included in the present study.

**Table 1.** Caries frequency for each species studied.

| Species | Observable teeth | Carious teeth | % caries |
| --- | --- | --- | --- |
| Early *Homo* | 44 | 2 | 4.55 |
| *P. robustus* | 318 | 7 | 2.20 |
| *H. naledi* | 147 | 2 | 1.36 |
| *A. sediba* | 16 | 0 | 0.00 |
| *A. africanus* | 328 | 0 | 0.00 |

**Table 2.** Hominin specimens with caries. Tooth: first letter, L (left), R (right); second letter, L (lower), U (upper); M1, first molar, P2, second premolar.

| Species | Specimen | Tooth | Position | Severity | Lesion # | Described |
| --- | --- | --- | --- | --- | --- | --- |
| *P. robustus* | SK 23 | LL M1 | Occlusal | 1 | 1 | This study |
| *P. robustus* | SK 23 | RL P2 | Occlusal | 1 | 1 | This study |
| *P. robustus* | SKX 5023 | RL M1 | Mesial/Distal | 2 | 3 | Grine et al. (1990) |
| *P. robustus* | SK 55 | LU M1 | Buccal | 1 | 2 | Robinson (1952) |
| *P. robustus* | SK 13/14 | LU M2 | Occlusal | 2 | 2 | Robinson (1952) |
| Early *Homo* | SK 15 | LL M1 | Mesial | 2 | 1 | Clement (1956) |
| Early *Homo* | SK 15 | RL M2 | Mesial | 2 | 1 | Clement (1956) |
| *P. robustus* | DNH 40 | LUM3 | Root/Mesial | 2 | 1 | Towle et al. (2019) |
| *H. naledi* | UW 101-001 | RL P2 | Distal | 2/3 | 1 | This study |
| *H. naledi* | UW 101-001 | RL M1 | Mesial | 2/3 | 1 | This study |
| *P. robustus* | SKW 5 | RL M1 | Occlusal | 1 | 1 | This study |
| *P. robustus* | SKW 5 | LL M1 | Occlusal | 1 | 1 | This study |

Two teeth belonging to the mandible SK 23 (*P. robustus*) have characteristics of caries on their occlusal surface. The left first molar and right second premolar have larger and darker colored fissures on their occlusal surface than do their antimeres (Figure 1a). However, due to post-mortem matrix in these depressions (Figure 1b), it was difficult to confirm whether these were carious through macroscopic observations. A micro-CT scan of these teeth supports the conclusion that this individual had carious lesions. Enamel under the depression on the occlusal surface of the right second premolar appears to have patches of less dense material, and a patchy track of less dense enamel extends towards the Enamel-Dentine Junction (EDJ; Figure 1d,e). This demineralized enamel near the occlusal surface fits a diagnosis of occlusal caries; the line towards the EDJ is likely also carious in nature, though it may have experience further postmortem cracking due to caries weakening the tissue. Therefore, the difference compared with its antimere (Figure 1a), large size of the depressions (Figure 1b), and lower density enamel underneath (Figure d,e), are all suggestive that a carious lesion had been active in the occlusal fissures of these teeth. These same features are also associated with the left first molar.

**Figure 1.**
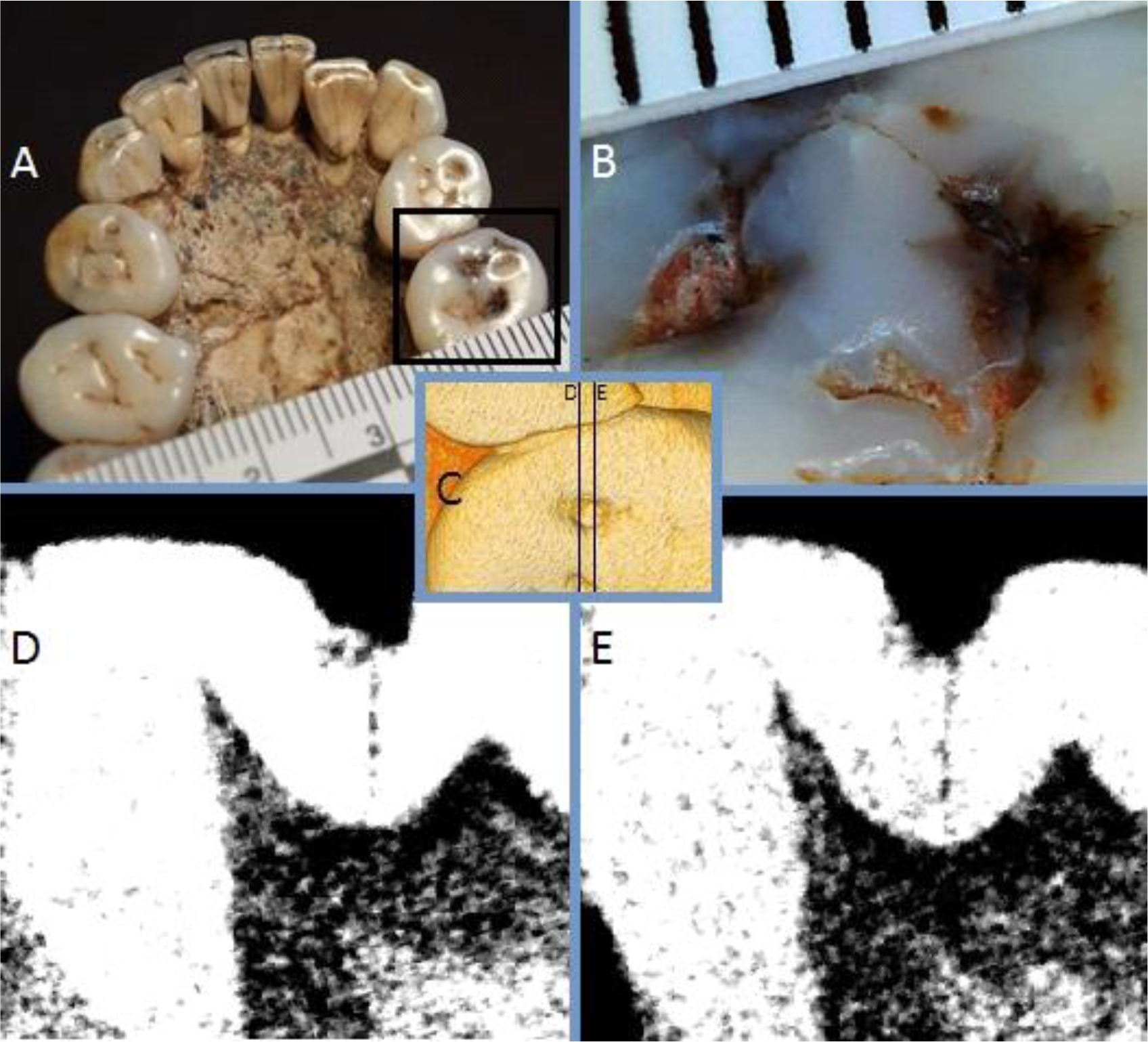
Caries in SK 23 (*P. robustus*). A) Occlusal view of mandibular teeth, with the right second premolar highlighted; B) Close-up of the occlusal surface of the right second premolar; C) CT reconstruction with the position of the two slices highlighted; D) CT slice toward the lingual part of the cavity; E) CT slice toward the buccal portion of the cavity.

Caries on *H. naledi* specimen UW 101-001 are the most severe of all South African hominins lesions yet described (Figure 2). It is clear that these lesions must have been active for several years since they had spread deep into the dentine. No reduction of crown wear is evident on this side of the mouth, so these lesions may not have affected normal mastication. The lesions formed on the interproximal area between the mandibular right second premolar and the adjacent first molar. The lesions likely started on the enamel of the interproximal surface, but has spread to effect the occlusal surface of the first molar as well as interproximal root surfaces. Due to postmortem sediment in the cavities it is not clear how deep into the dentine these lesions reach, or if the pulp chamber had been exposed. A micro-CT scan of this specimen was not available for study, but would likely give insight into the severity and spread of these carious lesions.

**Figure 2.**
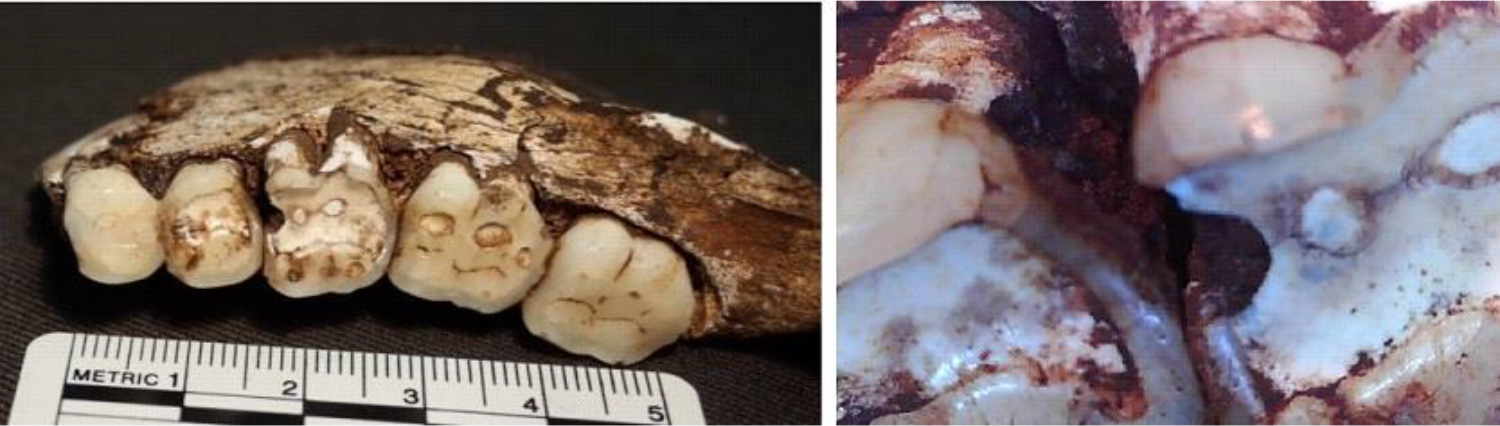
Carious lesions on the mandibular right second premolar (distal) and first molar (mesial). *Homo naledi* (UW 101-001).

The first molars of SKW 5 (*P. robustus*) have occlusal caries, based on the depth, coloration and pattern within the fissures (Figure 3a,b). When a micro-CT slice of this area is viewed, demineralization is evident deep into the enamel, and shows a pattern similar to enamel caries in clinical examples (Figure 3d). Pitting enamel hypoplasia is visible in the adjacent teeth, suggesting this condition may have facilitated or played a role in caries formation in this individual.

**Figure 3.**
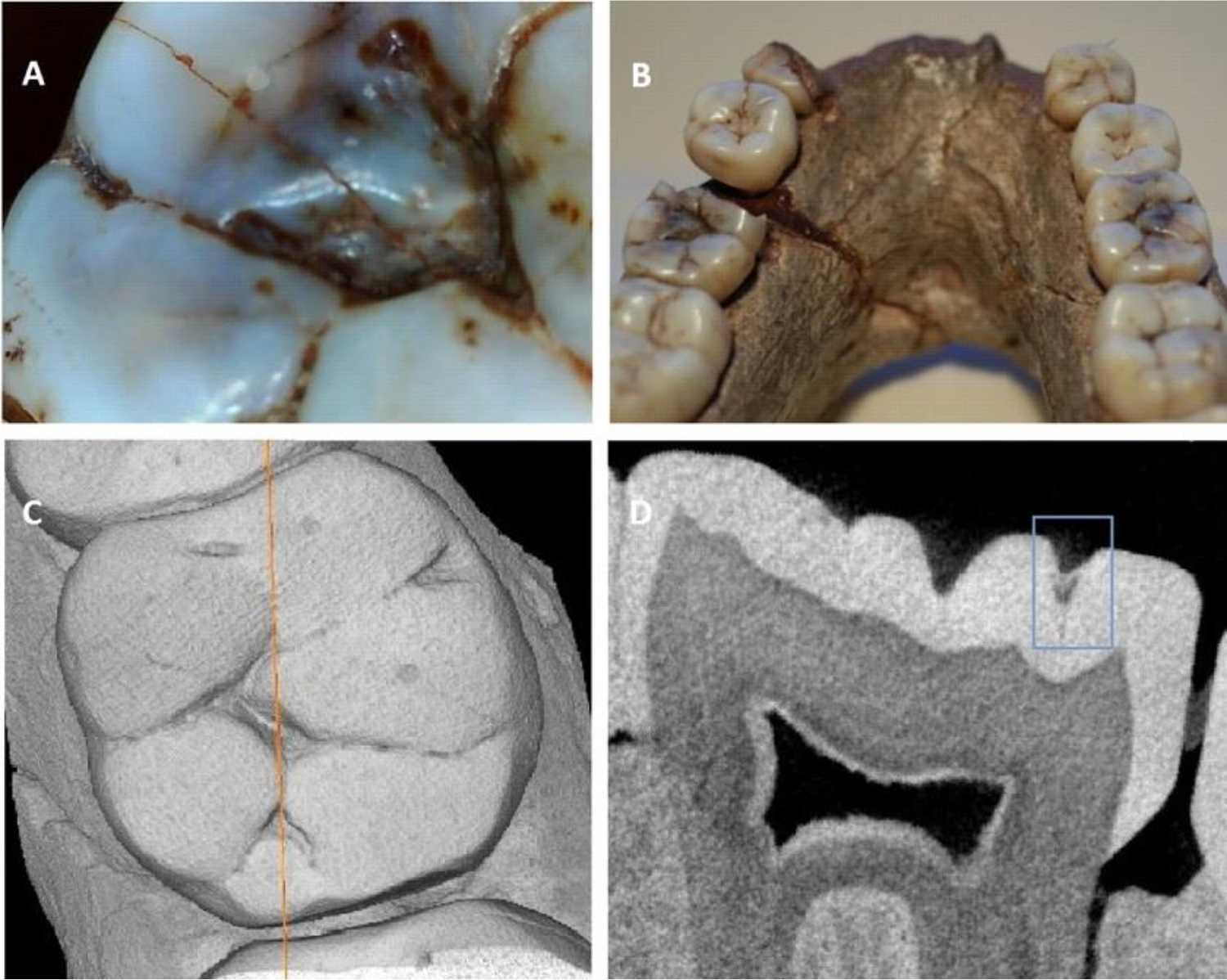
Caries on the occlusal surface of SKW 5 (*P. robustus*). A) Occlusal view of the right mandibular first molar; B) mandible displaying the two first molars with likely caries; C) CT reconstruction of the right first molar showing the position of the slice in D; D) CT slice of the right first molar, highlighting the area of demineralized enamel (blue square).

## Discussion

Carious teeth are likely to be lost or damaged post-mortem because they are more fragile than non-carious teeth. In support of this statement, all newly reported carious teeth were retained in a mandible or maxilla. Therefore, it seems likely that caries would have been more common in fossil hominins than the results of this study suggest. Recent research has highlighted that carious lesions may form more frequently on different dental tissues, or certain positions on a tooth crown, due to the presence of certain bacteria, pathologies, and dietary components (Meinl et al., 2010; Bignozzi et al., 2014; Novak, 2015; Takahashi & Nyvad, 2016). That said, all carious lesions ultimately share an etiology that is based on the presence of certain cariogenic bacteria and fermentable carbohydrates (Clarkson et al., 1987). Clearly, therefore, given the results of this and other recent studies (Grine et al., 1990; Trinkaus et al., 2000; Lanfranco & Eggers, 2012; Lacy et al., 2012; Humphrey et al., 2014; Lacy, 2014; Liu et al., 2015; Arnaud et al., 2016; Margvelashvili et al., 2016; Towle, 2019), cariogenic bacteria were prevalent in many, if not all, hominin species.

The frequency of caries on different tooth surfaces has predominantly been linked to the extent and speed of attrition on the occlusal surface. High rates of occlusal caries are associated with low attrition, whereas high levels of interproximal lesions are linked with high attrition (Hillson, 2008). However, high rates of interproximal caries may also be related to levels of calculus, in which plaque buildup in these areas may facilitate lesion formation (Tomczyk et al., 2013). There does not seem to be any evidence for calculus associated with caries in the South African hominins, although postmortem loss may have occurred. Enamel hypoplasia may also influence the formation of carious lesions. Defects may act as a site in which lesions can develop, but also may speed up the progression of a lesion, as the enamel may be more vulnerable to acid solubility (Hong et al., 2009; Rohnbogner & Lewis, 2016). Therefore, enamel hypoplasia seems likely associated with caries in *P. robustus*, with specimens SK 55, SK 13/14 and SKW 5 displaying both caries and enamel hypoplasia (Towle and Irish, 2019). Other pathologies and wear can also create an environment in which caries is more likely to form. For example, caries may develop as a response to unusual occlusion (Calcagno & Gibson, 1991). In populations with a moderate amount of caries, presumably such occurrences will be proportionally more likely to be an influencing factor of cavity formation.

In human agricultural groups maxillary teeth tend to display higher rates of caries than mandibular teeth (Lunt, 1974; Caglar et al., 2007; Esclassan et al., 2009; Novak, 2015). It is also more common for caries to be found on posterior teeth than anterior (Varrela, 1991; Slaus et al., 1997; Watt et al., 1997; Vodanović et al., 2005; Novak, 2015). Similarly, the positions most affected are interproximal and occlusal surfaces (Varrela, 1991; Srejic, 2001; Caglar et al., 2007; Esclassan et al., 2009; Slaus et al., 2011; Vodanovic et al., 2012). These differences in location likely reflect the greater complexity of posterior teeth and their larger size, in which more susceptible areas are present for caries to form (Hillson, 2001; 1996). For previously reported carious lesions in fossil hominins, the majority are located on interproximal areas of the tooth crown rather than root surfaces. This is likely due to periodontal attachment loss being uncommon in earlier hominins (Stamm et al., 1990; Bignozzi et al., 2014). However, root caries has been shown to be formed commonly by particular bacteria in modern humans (Shen et al., 2005) and may also develop in different acidic conditions than enamel caries (Shen et al., 2004). Fossil hominins in general tend to display interproximal carious lesions, this likely reflects the heavy occlusal wear in these populations, in which fissures and pits on the surface are quickly worn away with wear progressing too fast for lesions to form. However, in this study we highlight occlusal caries in *P. robustus*, and a root carious lesions has also recently been described in a *P. robustus* individual from the site of Drimolen (Towle et al., 2019). Therefore, clearly a variety of types of carious lesions are present in the fossil record, and may reflect dietary differences between samples.

Many human samples within the last 50,000 years show caries frequencies of around the same or below some of the species in this study (Kelley et al., 1991; Larsen et al., 1991; Lacy, 2014). With an agricultural diet, this figure varies more dramatically, and some populations had vastly greater rates. However, rather than a steady increase, as has been suggested, the rate is rather stable over the last two million years. Therefore, the commonly perceived notion that caries is a new disease, in which only recent populations of the genus *Homo* are affected, may be misleading (Brothwell, 1963; Armelagos & Cohen, 1984; Hildebolt & Molnar, 1991; Tillier et al., 1995; Lanfranco & Eggers, 2012; Guatelli-Steinberg, 2016). Instead it seems most hominin populations had a moderate rate of caries, typically 1-5% of teeth affected, and if certain groups fall either side of this range then there is usually a specific dietary or behavioral explanation. For example, this is the case for many agricultural diets (e.g., Varrela, 1991; Srejic, 2001; Vodanović et al., 2005; Slaus et al., 2011; Novak, 2015), hunter gather groups reliant on a particular cariogenic food (e.g., Humphrey et al., 2014), and a diet high in marine foods/terrestrial meat (e.g., Kelley et al., 1991; Larsen et al., 1991; Lacy, 2014). All major samples of fossils assigned to the genus *Homo*, and studied for the presence of dental pathologies, show carious lesions (e.g., Lanfranco & Eggers, 2012; Lacy et al., 2012; Lacy, 2014; Liu et al., 2015; Margvelashvili et al., 2016; Arnaud et al., 2016), and it looks like most of these samples will fall within the 1-5% range. From hominin species that are represented by large sample sizes and have been comprehensibly studied for caries, the only one to currently fall outside this 1-5% range is *A. africanus*.

The lack of caries in *A. africanus* potentially requires a dietary explanation. It seems unlikely a lack of carious bacteria is on its own responsible, particularly given the frequent occurrence of caries in other fossil hominins and extant great apes. Instead, some foods can actively restrict caries formation and are associated with low levels of lesions. In particular, tough, hard and fibrous foods can all create a less acidic oral environment due to high concentrations of saliva circulation (Moynihan, 2000; Prowse et al., 2008; Rohnbogner & Lewis, 2016). Meat is also generally thought to be associated with low rates of caries (Novak, 2015). Grit incorporated into the diet can create heavy wear and may therefore also mean caries lesions are less likely to form. Baboons masticate a significant amount of grit but also have an omnivorous diet containing a lot of tough foods such as meat, leaves and roots, likely explaining thier low levels of caries (Duray, 1992; Moynihan, 2000; Nystrom et al., 2004; Towle, 2017). Therefore, *A. africanus* may have also had an omnivorous diet. Occlusal wear and dental chipping frequencies for baboons and *A. africanus* are also very similar (Towle et al., 2017). A varied diet with lots of tough foods also fits with diverse results for *A. africanus* in isotopic, microwear, macrowear and biomechanical analysis (Sponheimer & Lee-Thorp, 1999; Van Der Merwe et al., 2003; Nystrom, 2004; Scott, 2005; Sponheimer et al., 2005, 2013; Strait et al., 2009; Towle et al., 2018). The results of the present study therefore adds support to the notion that there were significant dietary differences between *A. africanus* and other hominins species, in particular specimens assigned to the genus *Homo* and *Paranthropus*.

An explanation for the high caries rate in *P. robustus* is also required. The fact that these individuals have relatively high levels of wear means that caries would be expected to be low, especially on occlusal surfaces (Maat & Van der Velde, 1987; Moynihan, 2000). Therefore, along with the presence of some severe lesions, cariogenic foods may have frequently been consumed, i.e. a similar proportion of such cariogenic foods in the diet of a population without high occlusal wear would likely display higher rates of caries. The size of their molars is likely also an important factor, with extremely large complex molars creating a large area on which lesions could develop. The most likely explanation for the high rate of caries in *P. robustus* is the consumption of cariogenic fruits or vegetation matter, although another possible component may be the consumption of honey (Moynihan, 2000). Although, pitting enamel hypoplasia has also clearly inflated the caries rate by yielding suitable sites for lesions to form, cariogenic food and bacteria would have still been a necessary prerequisite for caries formation.

The severe lesions on teeth belonging to *H. naeldi* and early *Homo* also suggests cariogenic foods were commonly consumed in these populations. Large lesions have also been described in specimens outside of present day South Africa, including belonging to *Homo neanderthalensis*, early *Homo* and Pleistocene *H. sapiens*. Therefore, clearly large carious lesions have been common throughout the history of the genus *Homo*, and may relate to cooking or the consumption of cariogenic tubers and fruit. Honey is commonly consumed in many recent hunter-gatherer groups, and may therefore also be a potential factor involved in caries formation (Pontzer et al., 2018). More broadly, caries has also been found to be relatively prevalent in other wild great apes, and a lesion has been described in a Middle Miocene hominid (Stoner, 1995; Miles & Grigson, 2003; Fuss et al., 2018). The relatively high rate of caries may therefore be broader and be a hominid characteristic, although clearly there is much variation between species which allows dietary and behavioral inferences for fossil samples.

## Conclusions

The results of this study show distinct species differences, which likely reflect dietary and food processing variances among hominin groups. Caries is not as uncommon as previously thought in non-agricultural hominin groups. In fact, the fossil hominins in this study had rates similar and, in some cases, higher than modern human groups both pre-and post-agriculture. The results add support to recent findings that hominin groups tend to have 1-5% of teeth with carious lesions, with groups above or below this threshold usually having a specific dietary or behavioral property explaining the low/high rate of caries. Overall, cariogenic foods have clearly been commonly consumed throughout human evolution and cariogenic bacteria have also been present, and caries frequency is therefore a useful tool for reconstructing diet of past populations.

## Acknowledgements

The authors thank L. Berger and B. Zipfel from the University of the Witwatersrand and S. Potze from the Ditsong Museum of South Africa for access to their collections. For producing the CT scans provided by Ditsong Museum of South Africa, we thank J.J. Hublin and the Department of Human Evolution, Max Planck Institute for Evolutionary Anthropology. For technical assistance, we thank M. Skinner. This research was supported by a studentship to the first author from Liverpool John Moores University.

